# Using imputed genotype data in the joint score tests for genetic association and gene-environment interactions in case-control studies

**DOI:** 10.1101/062075

**Authors:** Minsun Song, William Wheeler, Neil E. Caporaso, Maria Teresa Landi, Nilanjan Chatterjee

## Abstract

**Background:** Genome-wide association studies (GWAS) are now routinely imputed for untyped SNPs based on various powerful statistical algorithms for imputation trained on reference datasets. The use of predicted allele count for imputed SNPs as the dosage variable is known to produce valid score test for genetic association.

**Methods:** In this paper, we investigate how to best handle imputed SNPs in various modern complex tests for genetic association incorporating gene-environment interactions. We focus on case-control association studies where inference in an underlying logistic regression model can be performed using alternative methods that rely on varying degree on an assumption of gene-environment independence in the underlying population. As increasingly large scale GWAS are being performed through consortia effort where it is preferable to share only summary-level information across studies, we also describe simple mechanisms for implementing score-tests based on standard meta-analysis of “one-step” maximum-likelihood estimates across studies.

**Results:** Applications of the methods in simulation studies and a dataset from genome-wide association study of lung cancer illustrate ability of the proposed methods to maintain type-I error rates for underlying testing procedures. For analysis of imputed SNPs, similar to typed SNPs, retrospective methods can lead to considerable efficiency gain for modeling of gene-environment interactions under the assumption of gene-environment independence.

**Conclusions:** Proposed methods allow valid analysis of imputed SNPs in case-control studies of gene-environment interaction using alternative strategies that had been earlier available only for genotyped SNPs.

## Introduction

Genome-wide association studies (GWAS) are now routinely imputed for untyped SNPs with powerful imputation algorithms [Howie et al., 2009; Browning and Browning, 2009; Li et al., 2010; O’connel et al., 2016; Loh et al., 2016] up to various reference panels such as the Hapmap (The International HapMap Consortium, 2005) and the 1000 Genomes (The 1000 Genomes Project Consortium, 2010; The 1000 Genomes Project Consortium, 2012; Sudmant, P. H. et al., 2015). Standard association tests for imputed SNPs are performed using the predicted allele count as the underlying dosage variable of the association model. Many earlier fine mapping studies based on the Hapmap panel have successfully used imputation for better characterization of common susceptibility SNPs within regions initially discovered through typed SNPs. More recently, imputation based on the 1000 Genome reference panel in existing GWAS for several traits have led to the discovery of new susceptibility loci containing uncommon or rare susceptibility variants [Guerreiro et al., 2013; Wang et al., 2014; Horikoshi et al., 2015].

The use of expected allele count for imputed SNPs as the dosage variable is known to produce valid score-test for genetic association [Marchini and Howie, 2010]. In this paper, we investigate how to best handle imputed SNPs in various modern complex tests for genetic association incorporating gene-environment interactions. In particular, we focus on case-control association studies where inference in an underlying logistic regression model can be performed using various alternative methods that rely on varying degree on an assumption of gene-environment independence in the underlying population. As increasingly large scale GWAS are being performed through consortia effort where it is preferable to share only summary-level information across studies, we also explore how these methods could be implemented in the context of meta-analysis. We study type-1 error and power of alternative methods using extensive simulation studies. An application of the methods is illustrated through a re-analysis of the National Cancer Institute GWAS of lung cancer that has been imputed by the 1000 Genome reference panel.

## Methods

### Options for Joint-Test of Association for Genotyped SNPs

We assume the main goal of our study is to test the association of disease-status (*D*) with genotype status ( *G*) of marker SNPs in the presence of a set of environmental risk factors ( *X*) that are known to be associated with the disease. We consider logistic regression to specify the disease-risk model in the form

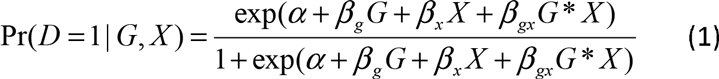

where an interaction term between *G* and *X* is incorporated to allow the effect of the genetic factor, as measured in the odds-ratio scale, to vary by the level of the environmental factors. Commonly, SNP genotypes ( *G*) are coded as allele count assuming a linear-trend model for association with the underlying trait. More generally, genotype could be coded according to dominant, recessive or a two degree-of-freedom saturated model. A joint-test for genetic association under the above model corresponds to a global null hypothesis in the form

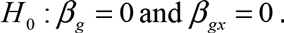

For genotyped SNPs, a multi degrees-of-freedom joint-test of association and interaction has been studied earlier [Kraft et al., 2007]. Typically, the analysis is performed based on standard prospective logistic regression analysis of case-control data.

Alternatively the analysis can be performed based on a retrospective-likelihood [Chatterjee and Carroll, 2005] that allows enhancement of power by exploitation of an assumption of gene-environment independence in the underlying population. Under gene-environment independence assumption, a case-only analysis can also be performed for inference on the logistic regression interaction parameter [Peigorsch et al., 1994], but it is not suitable for joint-testing of genetic association and interaction. The use of gene-environment independence assumption, however, can lead to serious bias in both the joint-and interaction-tests when the underlying assumption of gene-environment independence is violated [Albert et al., 2001;Mukherjee et al., 2008; Mukherjee et al., 2012].

A third alternative for joint-testing of association and interaction is to use an empitical-Bayes type inferential procedure that allows data adaptive shrinkage between estimates obtained from the prospective and retrospective likelihoods to strike a balance between efficiency and bias incurred by gene-environment independence assumption. Extensive simulation studies have shown that methods that exploit gene-environment independence assumption, such as retrospective-or EB- method, have substantial potential to improve power for gene-environment case-control studies compared to standrd prospective logistic regression [Mukherjee et al., 2008; Mukherjee et al., 2012]. The risk of false positives due to gene-environment correlation is generally low in many realistic situations and can be further minimized using the data adaptive EB or various types of two-stage procedures [Cornelis et al., 2012; Mukherjee et al., 2012].

### Derivation of Score-Tests

A major advantage of score-test, compared to Wald-or Likelihood-ratio test (LRT), is that it only requires imputation under the null model of no association and thus can easily incorporate expected dosage returned by popular imputation algorithms. Further for the analysis of less common and rare variants, score-tests may have more robust properties than Wald test or LRT as the number of cases or/and controls can be sparse in variant genotype categories.

Suppose that data consist of (*D_u_, X_u_, G_u_*), *u* = 1,…,*n* where *D_u_*, *X_u_*, and *G_u_*, respectively, denote the disease status, environmental exposure, and SNP-genotype status for subject *u*. Let *Z* = (1, *X*) and *W* = (*G, G* * *X*) denote a partitioning of the design matrix associated with the “nuisance” parameters, *η* = (*α,β_x_*), and the parameters of interest, *θ* = (*β_g_,β_gx_*), respectively, for the underlying logistic regression model.

### Prospective (PT) Method

The standard prospective likelihood of case-control data is derived as

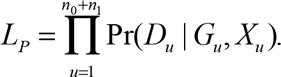

Under the prospective-likelihood, the score-function for *θ* is given by

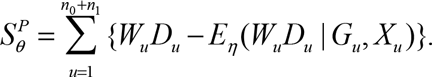

Under the null hypothesis of no association,

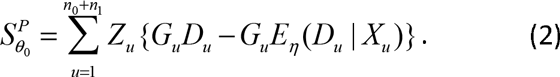

The maximum likelihood estimator (MLE) of the nuisance parameters *n* under the null model can be estimated by fitting the null model

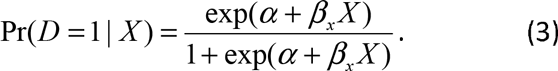

The multivariate score-test-statistic can be computed as

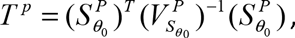

where 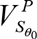 is the variance-covariance matrix for the score-vector accounting for uncertainty associated with estimation of the nuisance parameters. One can estimate 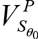 using the efficient information matrix in a model-based fashion or using the empirical variance-covariance matrix of the associated influence function to achieve robustness against mis-specification of the null model.

### Retrospective (RT) Method

The retrospective likelihood for case-control data is given by

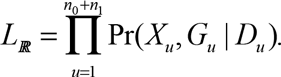

It has been long known that inference for the parameters of interest under underlying logistic regression model is equivalent under the retrospective and prospective likelihoods for case-control data when no assumption is made about joint distribution of the underlying risk-factors, i.e. *G* and *X* in our example [Prentice and Pyke, 1979]. However, if an assumption of gene-environment environment independence is invoked, then more efficient inference is possible under the retrospective likelihood. In particular, Chatterjee and Carroll [2005] have previously shown that under the assumption of gene-environment independence, but without any further restriction on the distribution of *X*, inference under retrospective likelihood can be made using a “profile-likelihood” of the form

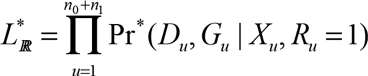

where the conditioning *R* = 1 is introduced to indicate the selection mechanisms of subjects into the sample under the case-control sampling scheme. Derivation of 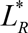 requires specification of population genotype frequencies, either using two parameters under a general multinomial model or using a single parameter under the assumption of Hardy Weinberg Equilibrium (HWE). Thus, for the retrospective likelihood, we expand the nuisance parameter vector as *η*^*^ =(*η*, *γ*) that the nuisance parameters include both parameters of the disease-risk and genotype frequency models.

The score-function for association parameters of interest for the retrospective likelihood can be derived as

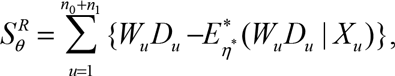

where 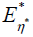 denotes expectation with respect to the joint probability distribution of *D* and *G* given *X* and *R* = 1.

Under the null,

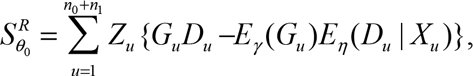

which differs from the corresponding score-vector (equation (2)) is derived under the prospective likelihood only in the way the expectation is derived in the second term. In particular, under the retrospective likelihood, the expectation term is evaluated under the assumption of gene-environment independence while the prospective likelihood does not require any such assumption. Under the null hypothesis, the parameters of the null model (3) can be estimated using standard prospective logistic analysis since the MLE under the retrospective-and prospective-likelihoods are the same as we allow the distrbution of non-genetic risk-factors to be completely unspecified. Further, under null, MLE associated with genotype frequency model *γ* can be obtained from the pooled sample of the cases and controls. The multivariate-score test can now be derived under the retrospective likelihood following the same-steps as that for described for the propsective likelihood (see Supplementary Methods Section 1.2 for complete details).

### EB Procedure

Implementation of the original EB procedures requires parameter estimates from the prospective-and retrospective-likelihood methods. The estimate itself cannot be directly derived from a likelihood and thus derivation of a score-test for this procedure is not straightforward. As an alternative, we propose a “score-type” test that could maintain some of the advatnages of the score-tests as described earlier and yet allow combining inference from the prospective-and retrospective-likelihoods in a data adaptive fashion to balance between bias and efficiency. We first note that any score-test can be written in the form of a Wald-like test-statistic

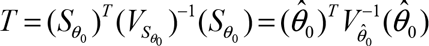
.

where 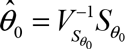 can be viewed as a one-step MLE starting from the null parameter value *θ*_0_=0. Thus, taking advantage of the above Wald-like representation of score-test, we propose an EB score-type test in the form

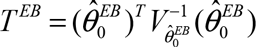

where the corresponding EB estimates and associated variance-covariance matrix are obtained by combining the one-step MLE estimates derived from the prospective-and retrospective-likelihood (see Supplementary Methods Section 1.3) using formulae analogous to those described for the original EB procedure [Mukherjee and Chatterjee, 2008].

Derivation of the PT, RT, and EB methods under a more general setting that allows accounting for additional covariates in the model is given in Supplementary Methods.

### Handling Imputed Genotype Data

Once the forms of the score-tests are derived with observed genotyped data, handling imputed genotype data for all the procedures is relatively straightforward as it simply involves replacing *G_u_* by *Ĝ_u_*, the expected value of genotype dosage taking into account predicted probabilities of different genotype values returned by the imputation algorithm. It is noteworthy how imputed genotype data are handled differentially in the prospective-and retrospective-score functions. Under the prospective-likelihood, the score-function for imputed genotype data takes the form

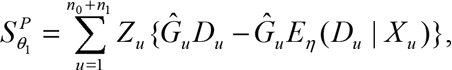

where the imputed genotype-dosage variable contributes to both terms of the left hand side of the equation. In contrast, under the retrospective-likelihood, the score function for imputed genotype data takes the form

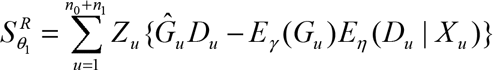

where the imputed genotype-dosage variable contributes only to the first term of the equation. The genotype frequency parameters, required in derivation of the retrospective-score function, can be estimated from imputed genotype data based on overall predicted genotype counts observed in the pooled sample of cases and controls. Derivations of efficient-information matrices and empirical variance-covariance matrices for the score-vectors follow the same steps as those for observed genotype data for each of the respective procedures. Finally, the derivation of the one-step MLEs and score-type test using the EB procedure follows the same steps as those described for observed genotype data.

## Analysis of NCI GWAS of Lung Cancer

We analyzed data from a GWAS of lung cancer generated at the National Cancer Institute. The dataset included 5713 cases and 5736 controls from four different study sites (Table 1). The samples were originally genotyped using a combination of Illumina GWAS platforms and were imputed using the 1000 Genomes Phase 2 reference panel using IMPUTE2 software [Howie et al., 2009]. The details of the studies can be found in several previous publications [Landi et al.,2009]. We evaluated the performance of the different methods in evaluating joint association of lung cancer with SNP-genotypes and genotype-by-smoking interactions. We derived score test under a logistic regression model where SNP genotypes were coded assuming additive effects. For modeling the effect of smoking status, recorded as current, former or never, we used two dummy variables. The resulting joint tests for association and interaction had three degrees of freedom. We also examined the two degree-of-freedom score-tests associated with only the interaction parameters of the model, but the underlying p-values were derived under the global null hypotheses of absence of both association and interactions. Figure 1 shows the quantile-quantile (Q-Q) plots for the interaction-only tests for the application of the PT, RT, and EB methods (left panel) and the joint-tests under the PT, RT, and EB methods (left panel) and the joint-tests under the PT, RT, and EB methods together with the test for main effect of *G* of the model without interaction (right panel), which were restricted to the analysis of ~5.3 million SNPs such that MAF > 0.05, the imputation quality reported to have info measure *I_A_* ≥ 0.5, and the p-values from all the seven tests are available. As the patterns are generally similar for the model-based and empirical variance estimators, we only show results using the former method. In general, the Q-Q plot associated with the interaction-only tests aligns close to the diagonal line indicating that all the methods are maintaining type-1 error well. The Q-Q plot neither shows any strong upward curvature near low p-values that could be indicative of the presence of many strong interactions in the data.

**Table 1.**
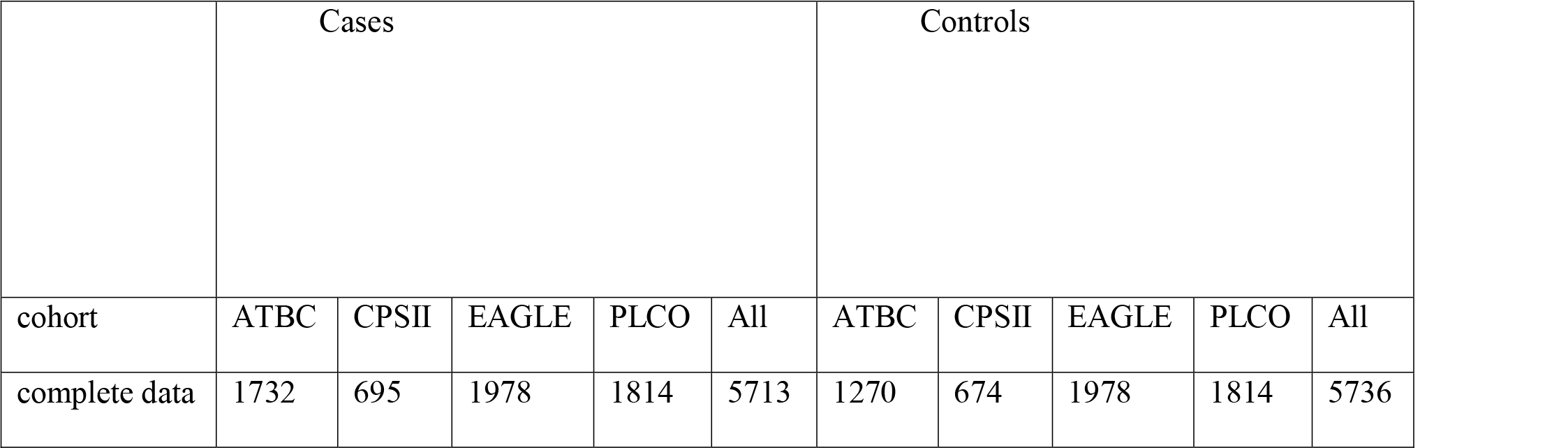
Distribution of cases and controls by cohorts in NCI GWAS.

**Figure 1.**
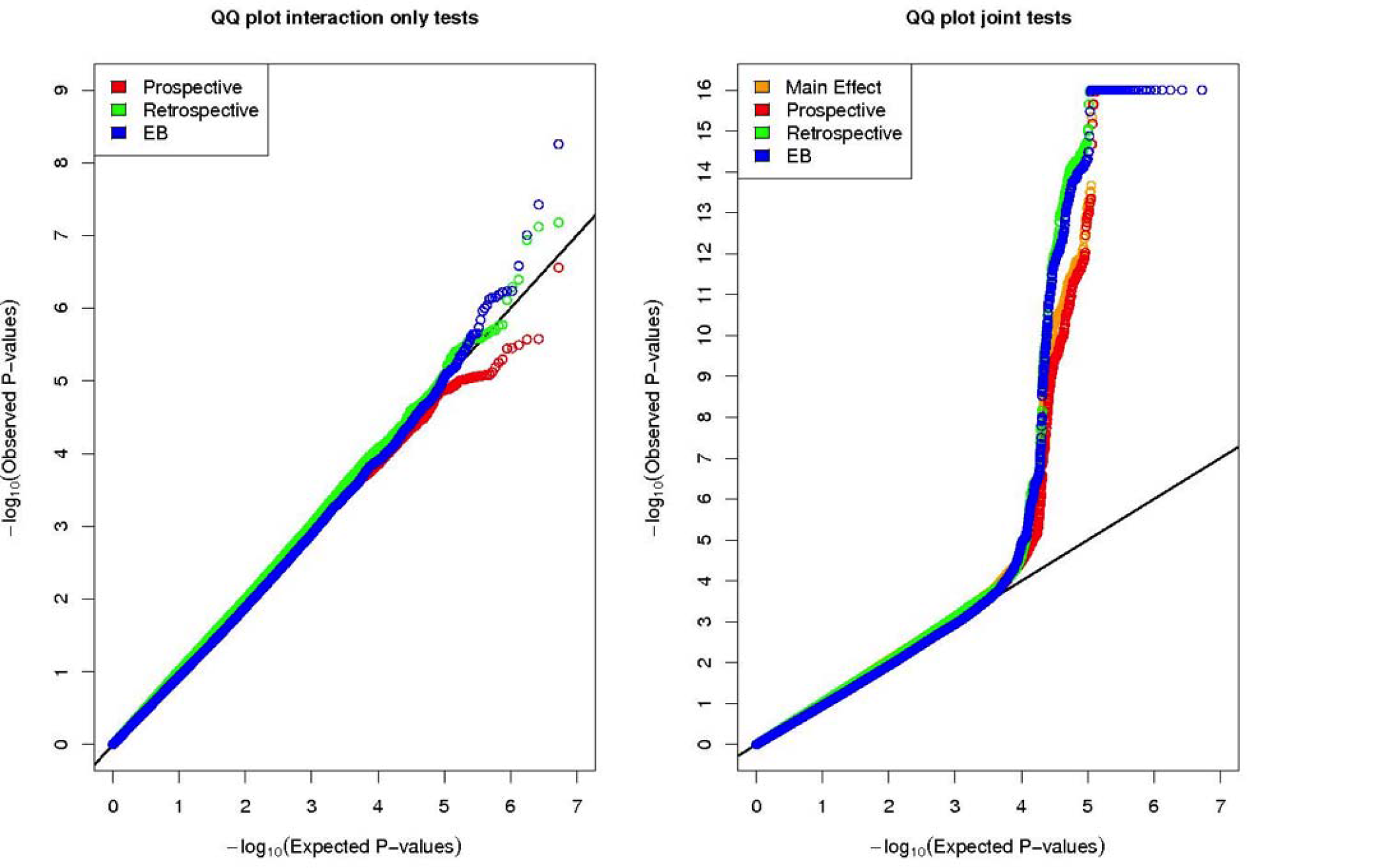
Quantile-quantile plots for the interaction-only, joint tests and tests for main effect of *G* in the analysis of National Cancer Institute Lung Cancer GWAS. Tests for associations are performed between risk of lung cancer and each of approximately 5.3 million common SNPs accounting for interactions with smoking status (never, former, and current) of the individuals. Each curve pertains to SNPs such that MAF > 0.05, the imputation quality reported to have info measure *I_A_* ≥ 0.5, and the p-values from all the seven tests are available. GWAS, genome-wide association studies; SNP, single nucleotide polymorphism; MAF, minor allele frequency.

In contrast, the Q-Q plot for the main-effect-only and joint tests of association and interaction clearly shows a strong upward curvature near the tail of the distribution. This pattern is largely driven by SNPs in the chromosome 15q25.1 region which are previously shown to be strongly associated with the risk of lung cancer (See Supplementary Figure S1 for plots after removal of this region). SNPs in this region, which contains multiple nicotine receptor genes, have been shown to be associated with both risk of lung cancer [Amos et al., 2008; Thorgeirsson et al., 2008] and smoking intensity [Thorgeirsson et al., 2008;Saccone et al.,2010]. However, no SNPs in this region has been reported to be associated with smoking status even in studies with extremely large samples size (N>100K) [The Tobacco and Genetics Consortium, 2010]. Thus it is interesting that in this region (x-axis p-value < 10^−4^), the RT and EB method, both of which exploit an assumption of independence of genotype and smoking status, consistently produce lower p-values for the SNPs than those from the main-effect only test and the joint-test under the PT method. It appears that, in this data, although gene-environment interactions by themselves are not identifiable at a high significance level, proper accounting for these effects using efficient methods are enhancing the detection of underlying signals captured by the joint test.

We also evaluated the performance of different methods including SNPs with lower MAF (MAF=0.01-0.05). In this setting, we observe that the Q-Q plots for the methods that used sandwich variance-covariance estimators were highly inflated indicating systematic problem with type-1 error rate control. The problem could be traced to small sample bias of the sandwich standard errors because of small number of non-smoking cases (N=355) who also carried variant genotype for rare SNPs in our study. When we combined non-smokers and former-smokers together to a single category, the bias went away (data not shown).

### Simulation Studies

We generated data on a binary environmental exposure variable which is assumed to follow Bernoulli (0.5) and be independently distributed of *G*. We simulated SNP genotype ( *G*) assuming HWE and minor allele frequency (MAF) value of 0.3 or 0.05. Given the values of *G* and *X*, we generated the binary disease outcomes for individuals from the logistic regression model (1). We chose *β_x_* = log(1.5) or *β_x_* = log(2) to allow the association of *X* with *D* to be either modest or strong, respectively. For evaluation of type-1 error, we assumed no genetic association, i.e. both *β_g_* = 0 and *β_gx_* = 0. For evaluation of power, we set *β_gx_* = log(1.2) and *β_g_* = log(1) or log(1.05). In all simulations *β_0_* was set such that an overall disease rate in the underlying population is about 5%. For evaluating type-1 error and power, we simulated 500,000 and 1,000 datasets, respectively, with each set consisting of 5,000 controls and 5,000 cases.

To evaluate validity and power of the methods when the SNP of interest may not be genotyped, we simulated haplotypes which consist of the SNP of interest and neighboring SNPs in linkage disequilibrium (Table 2). In one setting (left panel), the variant of interest was common (MAF=0.3) and could be imputed with high accuracy ( *R*^2^ =0.8) based on genotypes of the neighboring SNPs (See Stram [2004] for *R*^2^). In the other setting (right panel), the variant of interest was less common (MAF= 0.05) and could be predicted with moderate accuracy (*R*^2^ = 0.5) based on the genotype status of the neighboring SNPs. Assuming HWE in the general population, multi-locus genotypes of individuals were generated from simulted haplotypes. We analytically evaluated the conditional probability for genotype at the SNP of interest for each configuration of genotypes at the neighboring SNPs. For simulating “imputed dosage” for the SNP of interest, we simulated the genotype data for the neigboring SNPs first and then assigned predicted genotype probabilities for the SNP of interest using the known conditional probabilities. For analysis of each simulated data, we pretended that only the predicted probabilities, and not the actual genotypes, were available at the SNP of interest.

**Table 2.**
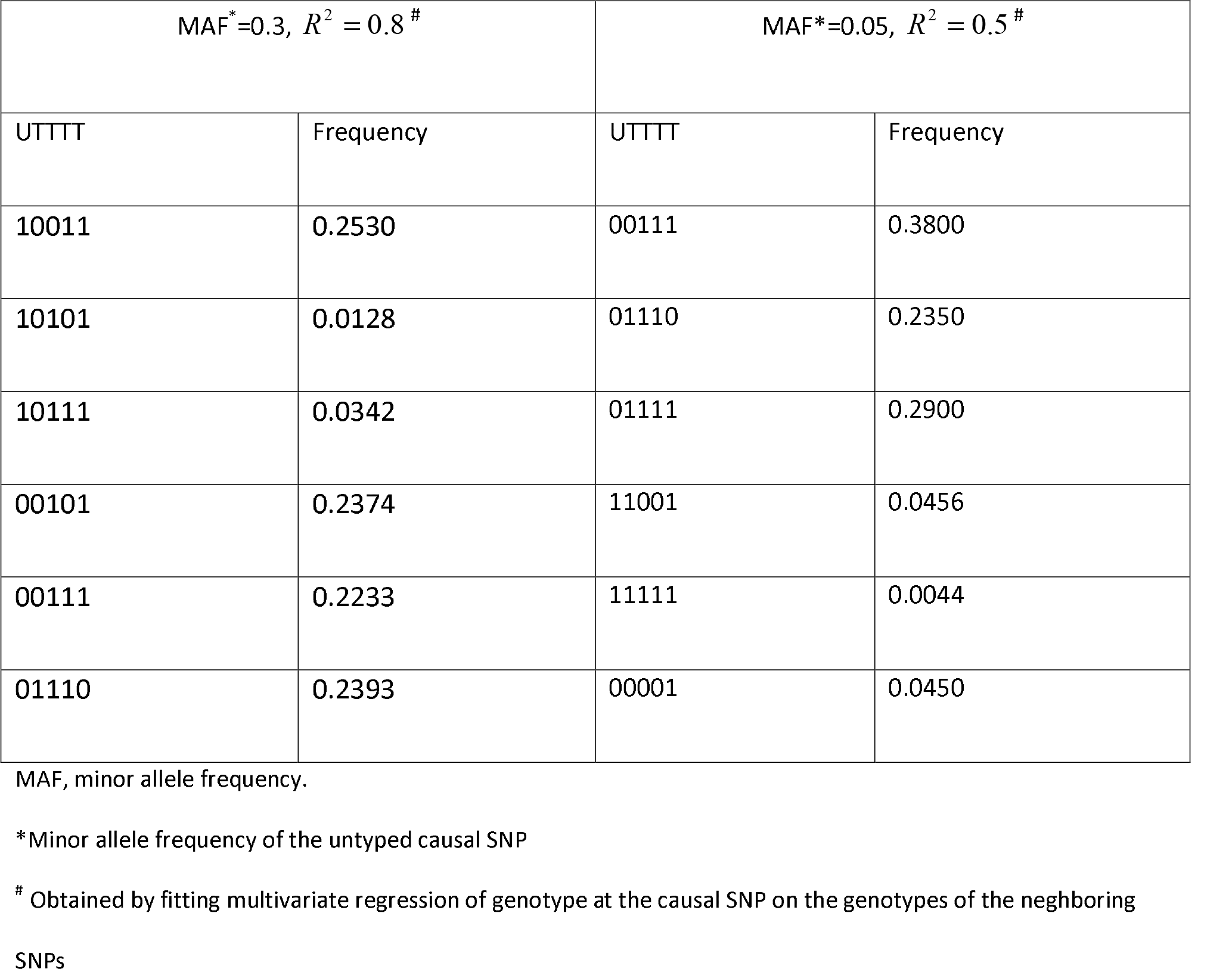
Haplotypes and their frequencies used for conducting simulation studies in scenario where underlying causal SNP is untyped and is assumed to be imputed based on neighboring genotyped SNPs. “U” and “T” indicate the untyped and typed SNP positions, respectively.

| MAF<sup>*</sup>=0.3, $R^2 = 0.8$ <sup>#</sup> | | MAF<sup>*</sup>=0.05, $R^2 = 0.5$ <sup>#</sup> | |
| --- | --- | --- | --- |
| UTTTT | Frequency | UTTTT | Frequency |
| 10011 | 0.2530 | 00111 | 0.3800 |
| 10101 | 0.0128 | 01110 | 0.2350 |
| 10111 | 0.0342 | 01111 | 0.2900 |
| 00101 | 0.2374 | 11001 | 0.0456 |
| 00111 | 0.2233 | 11111 | 0.0044 |
| 01110 | 0.2393 | 00001 | 0.0450 |

MAF, minor allele frequency.
\*Minor allele frequency of the untyped causal SNP
<sup>#</sup> Obtained by fitting multivariate regression of genotype at the causal SNP on the genotypes of the neighboring SNPs

We implemented each of the PT, RT, and EB score tests using either a model-based or an empirical variance-covariance estimator. However, in evaluation of the EB procedure, due to the lack of a model-based formula, the covariance between prospective and retrospective estimators was always evaluated based on empirical covariance of the underlying influenced functions. For each method, we evaluated the performance of both the joint-and interaction-only tests. In general, simulation studies show that the proposed methods perform well in maintaining type-1 error both at modertate (*α* = 0.05) and stringent (*α* = 0.0001) significance levels (Figure 2). In some scenarios, the RT method, when implemented with the sandwich variance estimator, showed a slight inflation over the nominal significance level. Across all the scenarios, the EB method was conservative, a pattern that has been reported earlier for analysis of typed SNPs and has been traced to the use of a conservative variance estimator [Mukherjee et al., 2012]. Employing the PT, RT, and EB methods on typed SNPs shows consistent results (Supplementary Figure S2).

**Figure 2.**
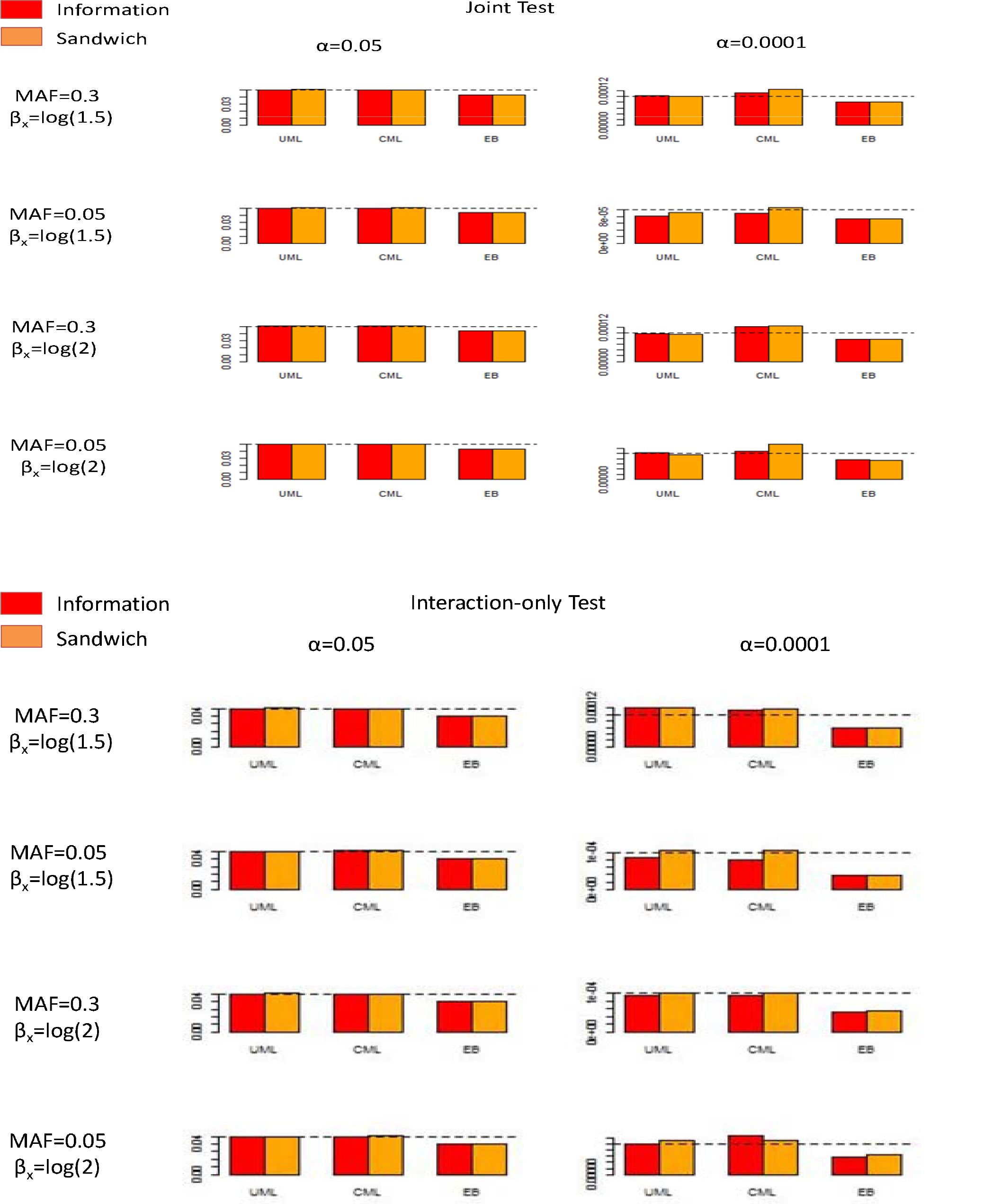
Simulation results for type-1 error for different procedures for testing untyped SNPs. The nominal significance levels are 0.05 and 0.0001. (top panels) Joint test, (bottom panels) Interaction-only test. Red and orange pertain to information-based variance estimator and sandwich variance estimator, respectively. MAF, minor allele frequency.

Simulation studies of power (Table 3) suggest that relative performance of three different methods was similar for untyped SNPs as has been reported for typed SNPs in earlier studies [Mukherjee et al., 2012]. In particular, the RT method had the maximum power, the PT method has the minimum power and the EB procedure performed in between. All methods lost power to a similar degree for the analysis of untyped SNPs compared to the analysis of the same SNP had it been typed. The use of model-based versus sandwich variance estimators did not have much effect in power for any of the methods (Supplementary Table S1).

**Table 3.** Simulation results for power of the joint-and interaction-tests for different procedures under various scenarios. For MAF=0.3, power is shown for nominal significance levels of 0.0001 and 0.001 for the joint-and interaction-tests, respectively. For MAF as 0.05, power is shown for the nominal significance level of 0.05 for both types of tests. In all settings, power was evaluated under an interaction odds-ratio=1.2. Results are shown when causal SNP is typed (bottom panels) or imputed (top panels). Variance is estimated based on information matrix.

| Untyped SNPs |  |  |  |  |  |  |  |  |
| --- | --- | --- | --- | --- | --- | --- | --- | --- |
| MAF | $\beta_x$ | $\beta_g$ | PT-joint | RT-joint | EB-joint | PT-int | RT-int | EB-int |
| 0.3 | log(1.5) | log(1) | 0.419 | 0.713 | 0.518 | 0.243 | 0.574 | 0.406 |
| 0.05 | log(1.5) | log(1) | 0.342 | 0.498 | 0.404 | 0.231 | 0.389 | 0.309 |
| 0.3 | log(1.5) | log(1.05) | 0.822 | 0.948 | 0.873 | 0.233 | 0.565 | 0.415 |
| 0.05 | log(1.5) | log(1.05) | 0.558 | 0.692 | 0.603 | 0.220 | 0.411 | 0.302 |
| 0.3 | log(2) | log(1) | 0.446 | 0.744 | 0.568 | 0.234 | 0.498 | 0.376 |
| 0.05 | log(2) | log(1) | 0.366 | 0.517 | 0.430 | 0.223 | 0.356 | 0.307 |
| 0.3 | log(2) | log(1.05) | 0.818 | 0.952 | 0.871 | 0.212 | 0.483 | 0.367 |
| 0.05 | log(2) | log(1.05) | 0.599 | 0.733 | 0.643 | 0.230 | 0.392 | 0.312 |
| Typed SNPs |  |  |  |  |  |  |  |  |

| MAF | $\beta_x$ | $\beta_g$ | PT-joint | RT-joint | EB-joint | PT-int | RT-int | EB-int |
| --- | --- | --- | --- | --- | --- | --- | --- | --- |
| 0.3 | log(1.5) | log(1) | 0.604 | 0.874 | 0.724 | 0.349 | 0.728 | 0.559 |
| 0.05 | log(1.5) | log(1) | 0.443 | 0.613 | 0.521 | 0.271 | 0.491 | 0.4 |
| 0.3 | log(1.5) | log(1.05) | 0.943 | 0.987 | 0.954 | 0.335 | 0.724 | 0.551 |
| 0.05 | log(1.5) | log(1.05) | 0.694 | 0.804 | 0.709 | 0.286 | 0.495 | 0.360 |
| 0.3 | log(2) | log(1) | 0.647 | 0.887 | 0.731 | 0.321 | 0.645 | 0.499 |
| 0.05 | log(2) | log(1) | 0.469 | 0.645 | 0.54 | 0.286 | 0.443 | 0.373 |
| 0.3 | log(2) | log(1.05) | 0.942 | 0.992 | 0.955 | 0.333 | 0.662 | 0.531 |
| 0.05 | log(2) | log(1.05) | 0.707 | 0.83 | 0.744 | 0.284 | 0.466 | 0.379 |

PT-joint, prospective joint test; RT – joint, retrospective joint test; EB-joint, empirical Bayes joint test; PT-int, prospective interaction test; RT-int, retrospective interaction test; EB-int, empirical Bayes interaction test; SNP, single nucleotide polymorphism; MAF, minor allele frequency.

## Discussion

In summary, we propose various types of score tests for genetic association and gene-environment interactions for analysis of case-control studies. Similar to standard tests for genetic association, in these methods, imputed genotype data for untyped SNP could be handled by simply substituting genotype values with predicted dosage that could be available from popular imputation software.

The prospective and retrospective score-tests are derived directly from the underlying likelihoods for case-control studies. We derived the score-test for the EB procedure using underlying one-step maximum-likelihood estimates of parameters obtained from the prospective-and retrospective-likelihoods. The one-step MLEs can also be used to perform multivariate meta-analysis of the parameters across studies and then derive various test-statistics based on meta-analyzed parameter estimates and their variance-covariance matrices. In our implementation of all the methods in the R software package CGEN (https://www.bioconductor.org/packages/release/bioc/html/CGEN.html) we allow returning of the one-step MLEs to facilitate meta-analysis.

Both analysis of simulated and real datasets suggest that the proposed methods can generally control type-1 error rates, but small sample bias could arise in the presence of sparse genotype-by-exposure cells, especially if sandwich variance estimators are used in some of these methods. Simulation studies of power show that the relative performances of the PT, RT, and EB procedure are quite similar for the analysis of untyped and typed SNPs. Although not studied directly, it can be anticipated that in the presence of gene-environment correlation in the population, the relative performance of these methods for their ability to control type-1 error would also be similar as has been reported in earlier studies [Mukherjee et al., 2012] for typed SNPs.

Although all the methods are valid for both continuous and categorical exposures, our numerical studies only involve categorical exposures. Future studies are needed to investigate performance of the proposed methods in the presence of continuous exposure and model mis-specification. It has been noted before that if the model for association of the disease with a continuous exposure is mis-specified, then the test for genetic associations and interaction could be biased due to underestimation of variance of target parameters under the mis-specified model [Tchetgen and Kraft, 2011]. We focus on test for genetic association and interactions using single genetic markers. Further studies are also merited how these methods could be extended for derivation of gene-level aggregate tests of genetic associations and interactions.

## Acknowledgement

Research of Minsun Song, Maria Teresa Landi, Neil Caporaso, and Nilanjan Chatterjee was supported by the intramural program of the Nationcal Cancer Institute.

## Software link

The proposed method has been implemented in open source software, available at https://www.bioconductor.org/packages/release/bioc/html/CGEN.html.

